# Intraspecific variability exceeds interspecific variability in thermal responses of Baltic Sea *Ostreococcus* (Mamiellophyceae) traits

**DOI:** 10.64898/2026.01.13.698997

**Authors:** Carina Peters, Marie Vogel, Luisa Listmann’, C.-Elisa Schaum’

**Author notes:** Corresponding author ‘Shared last author.

## Abstract

Trait variation within and between phytoplankton species as well as within regions, can shape community dynamics and ecosystem processes. A better understanding of this variability will improve the prediction of future ocean scenarios and the possible resilience of different phytoplankton groups. We investigated thermal response variation within and among three *Ostreococcus* species—a prominent picophytoplankton genus—using 31 strains from the Baltic Sea area and eastern North Sea. Across a temperature gradient ranging from 18°C up to 32°C, we quantified growth, cell size, and a proxy for chlorophyll *a* (Chl*a*) and characterized their thermal responses using modeling approaches. We further assessed how these traits vary with temperature and quantified interspecific, intraspecific and intraregional variability. Thermal performance traits showed high variability within species and regions and moderate variability between species. *Ostreococcus tauri* showed significantly higher thermal optimum and a trend for higher peak rates than *Ostreococcus mediterraneus*. Thermal safety margin and the sensitivity of growth to temperature varied substantially within all species. For size and Chl*a*, the species showed distinct temperature responses: *Ostreococcus mediterraneus* exhibited highest intraspecific variability and delayed responses at colder temperatures, *Ostreococcus tauri* responded linearly, with the *Ostreococcus mediterraneus* variant showing intermediate patterns. High intraspecific variation in *Ostreococcus* indicates substantial evolutionary potential, while interspecific differences may contribute to ecosystem stability under changing temperatures. This is especially relevant with predicted temperature-induced community shifts favoring smaller phytoplankton groups, highlighting the need to incorporate trait variability in ecosystem models.

## Introduction

Quantifying within and between species variability of picophytoplankton across different temperatures is critical for understanding their capacity to respond and adapt to climate warming. Intra- and interspecific variation of traits can significantly impact the functioning of ecosystems, community dynamics and composition (Andersson et al., 2022; Bestion et al., 2021; P. K. Thomas et al., 2024; Xiao et al., 2017). Temperature is a key environmental driver that affects metabolism, cell composition, productivity, and ultimately the carbon cycle (Anderson et al., 2022; Barton et al., 2018; Fernández-González & Marañón, 2021; Strock & Menden-Deuer, 2021; Toseland et al., 2013). While interspecific variation in the thermal adaptation of traits—especially among phylogenetically diverse taxa—has been documented (Anderson et al., 2021; Barton et al., 2023), the magnitude of intraspecific variation and variation among closely related species remains equally important to understand. Estimates of trait variability may also depend on the environmental and trait range over which they are assessed, as broader gradients can reveal patterns not captured by more limited measurements.

Picophytoplankton play a central ecological role in aquatic systems and will become increasingly important as warming shifts community composition toward smaller phytoplankton groups (Pulina et al., 2016; Suikkanen et al., 2013; Trombetta et al., 2019). Picoeukaryotes are generally less abundant than autotrophic prokaryotes, but no less relevant as they can occupy niches, e.g. through light or nutrient limited niche partitioning, (Agawin et al., 2000; Botebol et al., 2017; Rodríguez et al., 2005). They are important contributors to the carbon cycle, both in oligotrophic systems (A. Z. Worden, 2006; Alexandra Z. Worden et al., 2004) and eutrophic systems (Alegria Zufia et al., 2021; Martens et al., 2024; Spilling et al., 2025; Tamm et al., 2018). Among them, the picoeukaryote *Ostreococcus* is widely distributed from the Baltic Sea to the North Sea, Atlantic and Pacific Ocean (Demir-Hilton et al., 2011; Listmann et al., 2023; Simmons et al., 2016; Alexandra Z. Worden et al., 2004). As the smallest eukaryotic organism (∼1 µm; Courties et al. (1994)), *Ostreococcus* has become a widely used model organism for laboratory studies.

Despite the availability of strains representing all known clades and species, most studies focus on *Ostreococcus tauri (O. tauri)*, often repeatedly using the same long-cultured strains from laboratory collections, e.g. the strain RCC4221/OTH95, first isolated in 1995 (Barton et al., 2023; Bestion et al., 2020; Caló et al., 2022; Derelle et al., 2017; Ishikawa et al., 2024; Kay et al., 2021; Kubota et al., 2024; Vacant et al., 2022). Such strains may have undergone selection and evolution due to the time spent in culture collection and the way they are maintained (Lakeman et al., 2009). Relying on long-cultured, single strains may therefore limit our ability to capture the natural diversity of their responses (Bishop et al., 2022; Boyd et al., 2018; Hattich et al., 2017).

Consistent with this concern, studies have demonstrated ecotype-, species- and strain-specific differences in *Ostreococcus* responses to environmental conditions or between physiological traits (Cardol et al., 2008; Listmann et al., 2021; Schaum et al., 2012). For example, Schaum et al. (2012) found that *Ostreococcus* strains can differ in their plastic response to CO_2_ enrichment, which were largely correlated with geographic region rather than clades. Similarly, phytoplankton temperature traits can also vary across smaller geographical scales (Hattich et al., 2017; Santelia et al., 2026). Together, this highlights the need to study multiple, recently isolated strains and species across and within regions. While interspecific variability can enhance community stability through complementarity and niche partitioning (Godoy et al., 2020; Vallina et al., 2017), intraspecific variability, especially within regions, could drive substantial phenotypic diversity, providing a buffer against environmental fluctuations (Kremp et al., 2012) and future changes (Ajani et al., 2021; Bishop et al., 2022). Such regional variability may be especially important in the Baltic Sea, which experiences strong environmental gradients, rapid warming (Naumann et al., 2019; Ptak et al., 2026; Snoeijs-Leijonmalm et al., 2017) and increasing heatwave frequencies and lengths (Frölicher et al., 2018; Lindenthal et al., 2024; Pinto et al., 2024). However, it remains unclear how thermal responses vary among closely related *Ostreococcus* species within this region and whether they occupy distinct or overlapping thermal niches.

Here, we investigated intra- and interspecific variation in thermal responses across 31 *Ostreococcus* strains isolated predominantly from the Southern Baltic Sea, as well as the Skagerrak and the North Sea. Specifically, we tested whether (i) thermal tolerance and growth responses vary within and between species, (ii) substantial variability exists within a single region (Kiel Basin), and (iii) patterns of variability are consistent across traits. Given the environmental variability of the Baltic Sea, we expected pronounced within-species and within-region variation, consistent with adaptive potential under future warming. Furthermore, due to their close relatedness and co-occurrence, we expect the studied species to exhibit at least partly overlapping thermal niches.

## Material and Methods

To assess thermal responses of traits, we measured growth rates and extracted cell size and a proxy for chlorophyll *a* content from cytometric measurements for 31 different *Ostreococcus* strains at temperatures from 18°C up to 32°C. This allowed us to compare thermal optima (T_opt_), the range of temperatures beyond T_opt_ at which growth is possible (thermal safety margin), the activation energy (E_a_; or thermal sensitivity, i.e. the slope leading up to T_opt_) and the maximum growth rate at T_opt_ (peak rate) of the different strains and species (Daniel Padfield et al., 2026). In addition, thermal performance curves (TPCs) and temperature dependent changes in size and chlorophyll *a* content revealed the level of phenotypic plasticity (ability to “produce distinct phenotypes in response to environmental variation”, Sommer (2020)) among strains and species, giving a more comprehensive picture about the responses to temperature and the variation therein.

### Isolation and stock culturing of Ostreococcus strains

21 of the *Ostreococcus* strains used in this study were isolated between March 2018 and March 2020 on different cruises in the Baltic Sea, in the Skagerrak and the North Sea (Fig. 1) as described in Listmann et al. (2021) and Listmann et al. (2023). Since then, 10 more strains have been isolated in March and April of 2021 from the Southern Baltic Sea (Bornholm Basin, Kiel Bight, Mecklenburg Bight) using the same method (see Table 1 for isolation time and precise isolation location). A strain here refers to a single cell isolate at a certain point and time.

**Figure 1.**
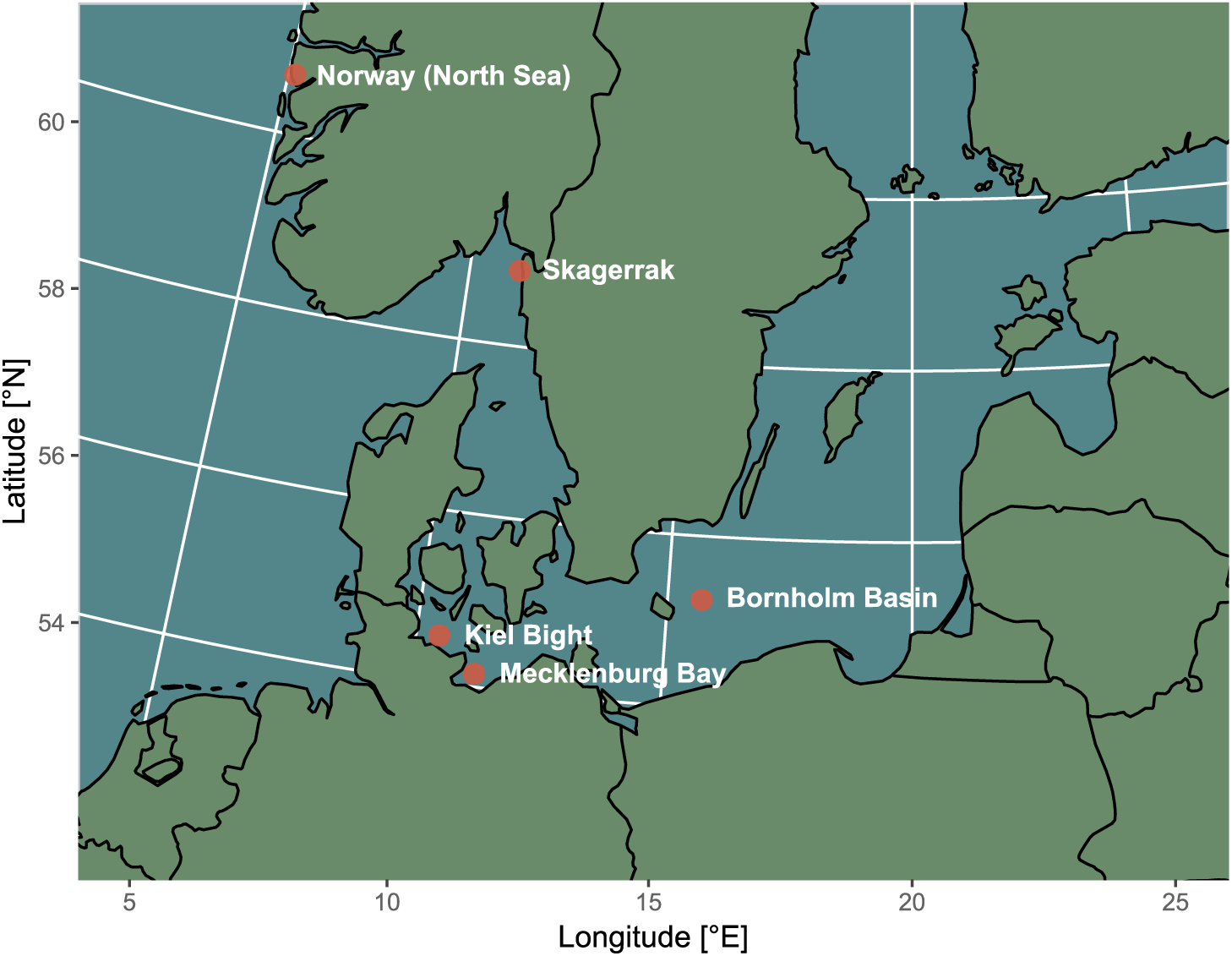
Map of the five main origin regions of *Ostreococcus* strains used in this study: three in the southern Baltic Sea (Kiel Bight, 17 strains; Mecklenburg Bay, 2 strains; Bornholm Basin, 6 strains), one in the Skagerrak (5 strains) and one in the North Sea near the coast of Norway (1 strain).

**Table 1.** Overview of the *Ostreococcus* strains as well as their origin region and time, species and location isolation coordinates. Strain names marked with an asterisk refer to strains isolated in Listmann et al. (2021) and Listmann et al. (2023), the remaining strains are the newer isolates.

| Strain | Region | Isolation Time | Temperature [°C] | Salinity | Species | Latitude | Longitude |
| --- | --- | --- | --- | --- | --- | --- | --- |
| AL505_St19.1* | Bornholm | March 2018 | 2.14 | 10 | *O. tauri* | 55.6119444 | 15.3038889 |
| AL505_St21.1* | Kiel | March 2018 | 1.76 | 15 | *O. tauri* | 54.5241667 | 11.3266667 |
| AL520_St5.13* | Kiel | March 2019 | 4 | 15 | *O. mediterraneus* | 54.6361111 | 11.0036111 |
| AL520_St2.29* | Kiel | March 2019 | 4 | 18 | *O. mediterraneus* variant | 54.3775 | 10.9136 |
| AL520_St2.30* | Kiel | March 2019 | 4 | 18 | *O. mediterraneus* variant | 54.3775 | 10.9136 |
| AL522_St03.A3* | Kiel | May 2019 | 11 | 18 | *O. mediterraneus* | 54.1986111 | 11.4688889 |
| AL522_St03.B5* | Kiel | May 2019 | 11 | 18 | *O. mediterraneus* | 54.1986111 | 11.4688889 |
| AL522_St03.C2* | Kiel | May 2019 | 11 | 18 | *O. mediterraneus* | 54.1986111 | 11.4688889 |
| AL522_St03.C5* | Kiel | May 2019 | 11 | 18 | *O. mediterraneus* | 54.1986111 | 11.4688889 |
| AL522_St03.F6* | Kiel | May 2019 | 11 | 18 | *O. mediterraneus* | 54.1986111 | 11.4688889 |
| AL524_St12.C5* | Bornholm | July 2019 | 19.5 | 8 | *O. mediterraneus* | 55.0502778 | 13.6666667 |
| AL530_St1.B4* | Kiel | October 2019 | 12 | 15 | *O. mediterraneus* variant | 54.6111111 | 10.3008333 |
| AL530_St1.B6* | Kiel | October 2019 | 12 | 15 | *O. mediterraneus* variant | 54.6111111 | 10.3008333 |
| AL530_St1.F2* | Kiel | October 2019 | 12 | 15 | *O. mediterraneus* variant | 54.6111111 | 10.3008333 |
| AL530_St10.D8* | Mecklenburg | October 2019 | 13.2 | 15 | *O. mediterraneus* variant | 54.3011111 | 11.3822222 |
| AL530_St11.B5* | Kiel | October 2019 | 13.2 | 15 | *O. mediterraneus* variant | 54.6727778 | 11.0347222 |
| Sweden_B2* | Skagerrak | March 2020 | 4 | 28 | *O. mediterraneus* | 58.879518 | 11.124977 |
| Sweden_B5* | Skagerrak | March 2020 | 4 | 28 | *O. mediterraneus* | 58.879518 | 11.124977 |
| Sweden_C8* | Skagerrak | March 2020 | 4 | 28 | *O. mediterraneus* | 58.879518 | 11.124977 |
| Sweden_F8* | Skagerrak | March 2020 | 4 | 28 | *O. mediterraneus* | 58.879518 | 11.124977 |
| Sweden_G8* | Skagerrak | March 2020 | 4 | 28 | *O. mediterraneus* | 58.879518 | 11.124977 |
| HE570_St150.D3 | Norway | March 2021 | 4 | 32 | *O. tauri* | 60.692 | 5.158 |
| AL551_St1.D2 | Kiel | March 2021 | 3 | 15 | *O. mediterraneus* variant | 54.7275 | 10.0797222 |
| AL551_St1.G2 | Kiel | March 2021 | 3 | 15 | *O. mediterraneus* variant | 54.7275 | 10.0797222 |
| AL551_St3.D5 | Kiel | March 2021 | 2.8 | 15 | *O. tauri* | 54.89 | 10.3566667 |
| AL551_St4.E9 | Kiel | March 2021 | 3 | 15 | *O. mediterraneus* variant | 54.4894444 | 10.5916667 |
| AL551_St7.F4 | Mecklenburg | March 2021 | 3 | 15 | *O. mediterraneus* variant | 54.4016667 | 11.8330556 |
| AL553_StBB23.E2 | Bornholm | April 2021 | 4.72 | 8 | *O. mediterraneus* | 55.2969444 | 15.7508333 |
| AL553_StBB23.E6 | Bornholm | April 2021 | 4.72 | 8 | *O. mediterraneus* | 55.2969444 | 15.7508333 |
| AL553_StBB29.B9 | Bornholm | April 2021 | 4.56 | 8 | *O. mediterraneus* | 55.1297222 | 16.0075 |
| AL553_StBB29.C11 | Bornholm | April 2021 | 4.56 | 8 | *O. mediterraneus* | 55.1297222 | 16.0075 |

All stock cultures of these strains are kept at their isolation location salinity at a common garden temperature of 18°C at nutrient replete conditions (Guillard’s F/2) on a 12:12 h light-dark cycle at 100-150 µmol m^−2^s^−1^ (Rodríguez et al., 2005) with regular batch transfers. Species identity was determined by sequencing the 18S rRNA gene and the internal transcribed spacer (ITS) region, spanning ITS-1, 5.8S rRNA, and ITS-2. Sanger sequencing of 18S rRNA forward sequences was done with EUROFINSGENOMICS® as described in Listmann et al. (2023). For the ITS-region the following primers were used, the reverse primer was used for sanger sequencing: forward primer 5′-GTAGGTGAACCTGCGGAAGGA-3′, reverse primer 5′-CCTTGGTCCGTGTTTCTAGAC-3′. Sequences were clipped and aligned using Codon Code Aligner (18S: 908bp; ITS: 1018 bp). 18S rRNA reference sequences of O*. tauri, Ostreococcus mediterrraneus* (*O. mediterraneus), Ostreococcus lucimarinus* and *Ostreococcus* clade B strain RCC809 were used for species identification (ostta12g00750, RCC2590_scf16, Olu_18S_fromgenome, Rcc809_18S_fromgenome respectively). We identified four *O. tauri* and 16 *O. mediterraneus* strains. However, 11 of the strains aligned to *O. tauri* in one part of the 18S rRNA sequence and to *O. mediterraneus* in the other part (Supplementary Fig. S1), thus matching neither of the two species. Comparison with other known *Ostreococcus* 18S sequences through the NCBIs Nucleotide BLAST (Zhang et al., 2000) also did not reveal a perfect match. ITS-region sequencing confirmed this group clusters separately from both our *O. tauri* and *O. mediterraneus* strains in a phylogenetic analysis (Supplementary Fig. S2) with standard bootstrapping (1000 iterations). This separation was further supported by unpublished whole-genome sequencing data (G. Piganeau, personal communication), which showed that although these strains share identical 18S rRNA gene sequences with *O. mediterraneus* via whole genome sequencing, they exhibit substantially higher genomic divergence, with >300,000 single nucleotide polymorphisms (SNPs) compared to <60,000 SNPs within *O. mediterraneus*. We have therefore grouped the 11 strains separately from the other *O. mediterraneus* strains and named them *O. mediterraneus* variant.

### Growth Curves

To obtain growth curves for all of our 31 different *Ostreococcus* strains, we exposed each of the strains to up to seven different temperatures (15°C, 18°C, 20°C, 22°C, 24°C, 26°C, 28°C) and some to an additional 30°C and 32°C if they did not show any sign of growth limitation at 28°C (see Supplementary Figure S3 for which strain was tested at which temperatures). Due to technical issues with the light at 15°C and 22°C, those cultures were excluded from final analysis. No data was collected below 15 °C for logistical reasons, as incubators capable of maintaining such low temperatures were not available at the time. The strains were cloned previous to the experiment through single cell picking on agarose plates between September 2022 and April 2023. Strains were kept in biological triplicates at 90-160 µE (light in incubators is not equal depending on position in relation to light) and nutrient saturated conditions (Guillard’s F/2) in an Infors HT (R) shaking incubator (60RPM) at a 12:12 h light to dark cycle. Culture positions were switched at random to account for the differences in light. Due to the large number of cultures (up to 31 strains in three technical triplicates per temperature, 926 experimental units in total), the experiment was conducted in three batches. Nine strains were grown at repeating temperatures as a control between batches (see Supplementary Table 1).

The cultures were acclimated to their respective temperature treatments for 8-12 generations (Padfield et al., 2016; Staehr & Birkeland, 2006) before they were transferred to the experimental phase, where cell counts were determined daily until a minimum of 9.5 generations (Eq. 2) or stationary phase was reached (Fig. 2a). For the acclimation phase one large master mix was inoculated to 3000 cells/ml with F/2 media of the according salinity and then divided up into three 20 ml technical replicates. For the experimental phase, each replicate was then inoculated individually to a starting concentration of 3000 cells/ml. This approach reduced the experimental variability to a minimum.

**Figure 2.**
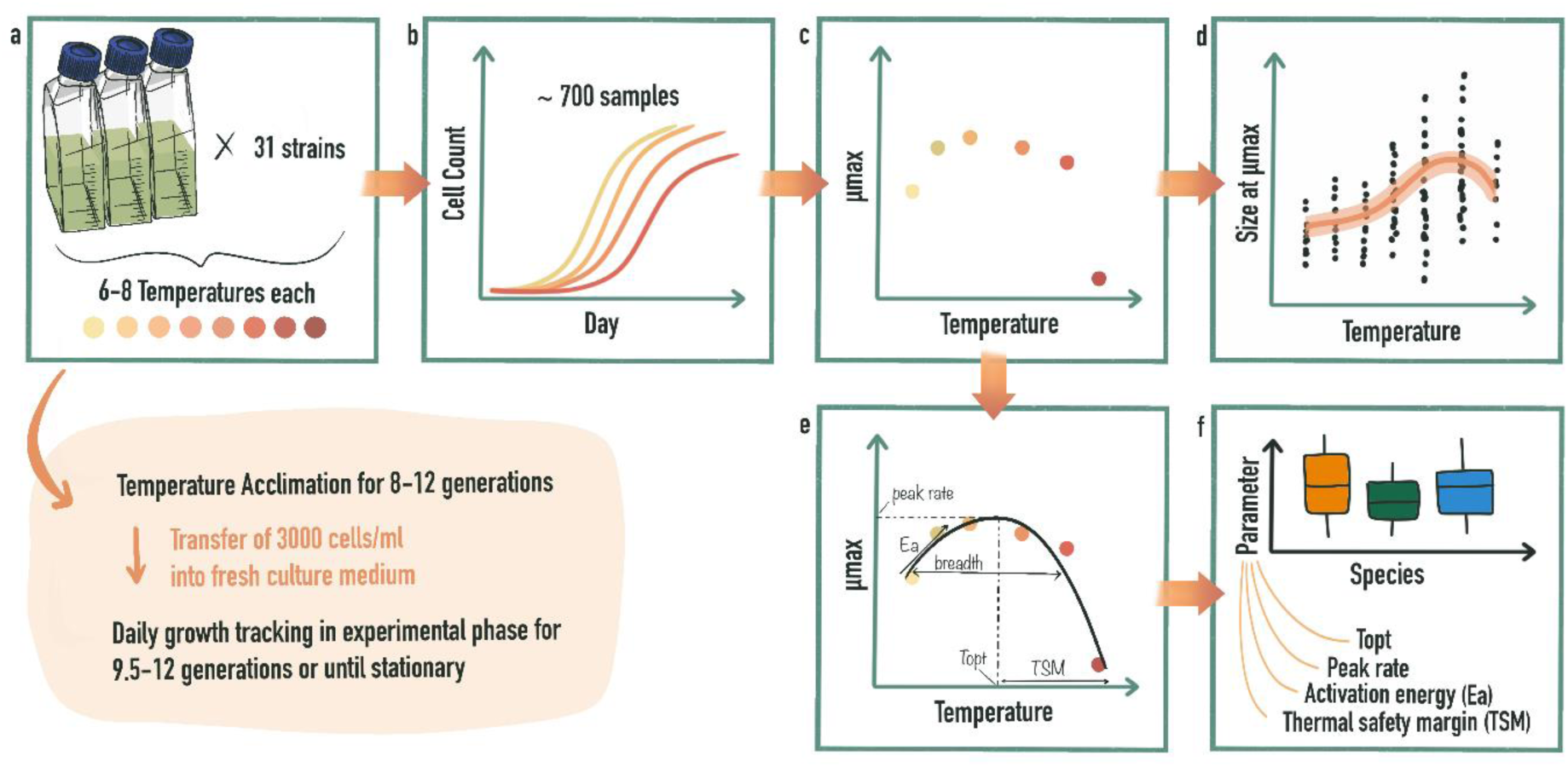
Illustration of method work-flow from a) experimental work to b) growth curve fitting and c) µmax extraction, which was then used for d) GAMM fitting of chlorophyll *a* content proxy and size at day of µmax and e) thermal performance curve fitting followed by f) thermal performance parameter extraction and comparison between species.

### Flow Cytometry

All cell counts were done via flow cytometry on the Beckman Coulter CytoFLEX using the FL3 Channel for chlorophyll *a* against size (FSC Channel). Since *Ostreococcus* only has a single chloroplast, one recorded signal can be approximately counted as a single cell (see Supplementary Fig. S4 for how populations were gated). A proxy for chlorophyll *a* content was extracted as mean relative fluorescence units (RFU) from the cytometric output for FL3 and used as a relative comparison between samples. For cell size, a standard curve with reference beads (Invitrogen™ Flow Cytometry Sub-micron Particle Size Reference Kit) was generated and then used to convert mean FSC-H measurements to cell size in µm.

### Growth rate and generation time calculations

For tracking the generation times during the experiment, the following formulas for growth rate and generations were used. Growth rate µ (d^−1^):

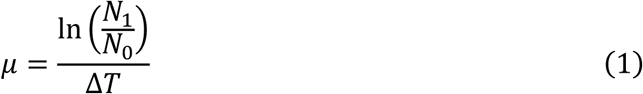

where N_1_ is the cell count at the point of measurement and N_0_ is the cell count at the start of the experiment. ΔT is the time interval in days between N_0_ and N_1_.

The number of generations g was calculated as follows, where ΔT is the time interval between sampling points in days, ln(2)/µ is the doubling time in days and µ is the growth rate d^−1^.

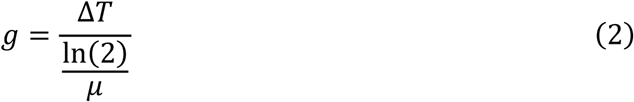

### Growth curve fitting and maximum growth rate extraction

All statistical analyses were done in the R programming environment (R version 4.5.0). Cell count data were log-transformed prior to any model fitting. For the extraction of µmax at each of the assay temperatures a non-linear least squares regression model was fitted using the nlsLoop (Version 1.0.0) package (Fig. 2b). Each replicate was fitted separately using the modified Gompertz model (gompertzm from the nlsMicrobio package Version 1.0-0) with up to 1000 attempts per curve. Initial parameter estimates (Supplementary Table 2) were chosen based on observed data. The estimated parameters, including µmax, were then extracted and used for further analysis (Fig. 2c). For estimation of the day when µmax was reached the following formula for inflection point calculation derived from the modified gompertz equation in the nlsMicrobio package (Supplementary Appendix A1) was used:

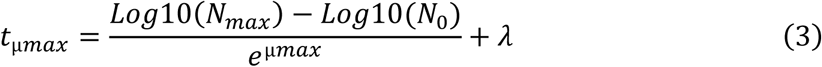

where t_µmax_ is the day µmax is reached (inflection point of the curve), µmax is the maximum growth rate, N_max_ is the carrying capacity, N_0_ is the starting concentration and λ is the lag phase. The resulting day was then rounded up or down to the nearest full day to extract the cytometric data of size and Chl*a* from that measuring day.

### Thermal performance curves

Thermal performance curves were fitted (Fig. 2e, Supplementary Fig. S3) to the maximum growth rate estimates using the nls.multstart (Version 2.0.0) and rTPC (Version 1.1.0) packages (Daniel Padfield et al., 2026; Padfield et al., 2021). To improve TPC fitting, zero values were added to the dataset for the strains that died in acclimation at their highest measurement temperature and therefore did not have a maximum growth—or any growth—estimation at that temperature. We acknowledge that this approach has limitations, but as growth rates often decline sharply beyond T_opt_, it can be challenging to identify temperatures at which strains exhibit minimal growth without experiencing mortality. One strain (AL551_St7.F4) was excluded for TPC fitting due to insufficient data points.

The available models of the rTPC package were reduced based on the *k* + 1 approach (Padfield et al., 2021), where *k* is the number of arguments used in the model formulation and + 1 meaning there should be at least one more data point than model arguments for a given model. The remaining models were tested on three representative strains (AL530_St11.B5, AL530_St10.D8 and AL522_St03.F6) spanning the upper and lower ranges of the dataset, including datasets with and without added zero values. Models that visually were not able to fit either one of those representative strains were excluded from further fitting. For the final round of model fitting the remaining models were fitted to all strains for further reduction based on the abilities to logically fit all 30 strains and estimate thermal performance parameters.

For the two final models (“lactin2_1995” and “tomlinsonphillips_2015”) the AICc (Akaike’s information criterion corrected for small samples size) values were calculated for best model selection per strain. The two chosen models are described as follows in equations for “lactin2_1995” (Eq. 4, Lactin et al. (1995)) and “tomlinsonphillips_2015” (Eq. 5, Tomlinson and Phillips (2015)).

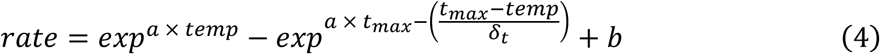

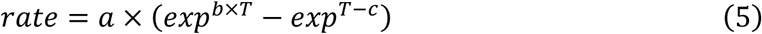

The thermal performance parameters (Fig. 2e) activation energy (E_a_), the maximum growth rate at T_opt_ (peak rate), thermal optimum (T_opt_) and thermal safety margin (TSM) were extracted per strain (Supplementary Table 3) using the calc_params function (Supplementary Table 4) in the rTPC package and compared across species (Fig. 2f). Confidence intervals (95%) for TPCs and parameter estimates were obtained by residual bootstrapping with 1000 replicates (Supplementary Table 3). Two strains (HE570_St150.D3 and Sweden_G8) yielded negative Ea estimates, indicating unreliable parameter identification from the available data. Because negative Ea values are not biologically interpretable within this framework, these estimates were treated as missing (NA), while all other parameters were retained. Three strains were excluded from all parameter comparisons as the data was either lacking a clear minimum at the lower end of the TPC (AL524_St12.C5, AL530_St10.D8) or no point beyond the highest measured temperature (AL553_StBB23.E2), and thus did not provide a reliable basis for parameter estimation.

For interspecific comparisons, parameters that violated parametric assumptions (peak rate, thermal safety margin) were analyzed using the Kruskal–Wallis test, whereas T_opt_ and E_a_ were analyzed using one-way linear models with species as a fixed factor. Model assumptions were tested with the Shapiro-Wilk-test and Levene’s test. Group means and confidence intervals were extracted with the emmeans package. Pairwise comparisons used Tukey’s HSD for linear models and Dunn’s tests for non-parametric analyses. Intraspecific variability was quantified as the coefficient of variation (CV; standard deviation/mean × 100). We note that the coefficient of variation (CV) should be interpreted with caution for *O. tauri*, as sample sizes for thermal performance parameters were low (n = 3–4).

### Species size and chlorophyll a content at µmax across temperatures (GAMM)

For analysis of species-specific differences in size and chlorophyll *a* content at µmax (Fig. 2d), we first compared cell size at µmax between species at a biologically relevant reference point, i.e. our common garden temperature of 18°C, at which all strains are routinely maintained. For comparisons of size a one-way ANOVA was conducted on strain-averaged log cell sizes and for the chlorophyll *a* content a non-parametric test (Kruskal-Wallis) was used, as data did not meet linear model assumptions. Pairwise comparisons were conducted with Tukey’s HSD and Dunn’s test, respectively.

To assess response of size and the chlorophyll *a* content proxy across temperature we each fitted a generalized additive mixed model (GAMM) for size and chlorophyll *a* content. Data was log-transformed prior to analysis. The model included species-specific smooth terms for temperature (k = 6) and a random intercept for strain identity using the REML method. Due to the limited number of temperature levels, k was restricted to 6. The coefficient of variation at each temperature was calculated as stated above.

## Results

### Intraspecific variability of thermal performance is higher than interspecific variability

The thermal performance curves (Fig. 3a, Supplementary Fig S3) showed variation within and between species in their curve shape and width. Thermal optima across all strains ranged from 22.41°C to 29.99°C and peak rate ranged from 0.78d^−1^ to 1.43 d^−1^. *O. tauri* displayed left skewed curves with moderate thermal sensitivity (E_a_ estimate: 0.18, CI: −0.001-0.36, Fig. 3c) towards T_opt_ and steep declines in growth rate beyond the thermal optimum. *O. mediterraneus* and the *O. mediterraneus* variant show a mix of left-skewed and more symmetrical bell-shaped curves, with similar thermal sensitivity (*O. mediterrraneus*: E_a_ estimate: 0.29, CI: 0.20-0.37, *O. mediterraneus* variant: E_a_ estimate: 0.28, CI:0.17-0.38).

**Figure 3.**
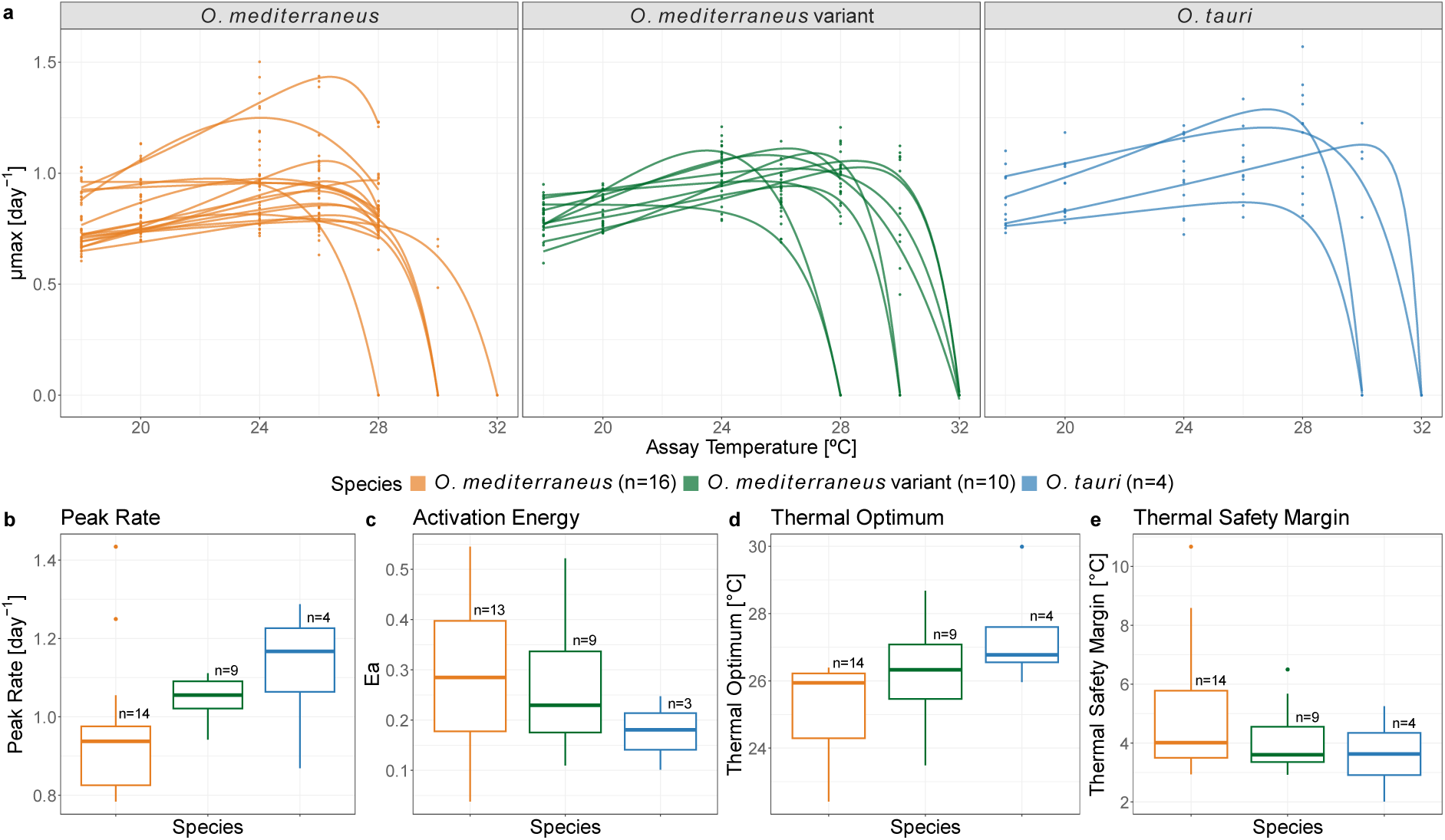
**a)** Fitted thermal performance curves of the different *Ostreococcus* strains used in this study, grouped by species. Predicted model fits of TPCs (lines) and extracted µmax data points (points) from growth curves are shown across assay temperatures. Boxplots of **b)** peak rate, **c)** activation energy, **d)** thermal optimum, **e)** thermal safety margin of the extracted parameters from the TPCs per strain are shown across species. The colors represent each species with orange for *O. mediterraneus*, green for the *O. mediterraneus* variant and blue for *O. tauri*. The boxes of the boxplots represent the 25th to 75th percentile of the data with the line within the box representing the median (50th percentile), the whiskers include the smallest and largest values without the outliers (outside 1.5 × IQR; shown as points).

There was a general trend for thermal performance parameters of *O. tauri* to be more different from *O. mediterraneus* than the *O. mediterraneus* variant, which often showed intermediate values. There was a significant species effect (ANOVA: F_(2, 24)_ = 3.675, p= 0.041) on T_opt_ (Fig. 3d), with a higher T_opt_ (Post-hoc Tukeys HSD: adj. p= 0.049) in *O. tauri* (mean 27.4°C ± 0.74 SE) compared to *O. mediterraneus* (25.3°C ± 0.40 SE). The *O. mediterraneus* variant exhibited an intermediate T_opt_ (26.4°C ± 0.49SE) relative to the two other species. Similarly, peak rate (Fig. 3b) differed significantly among species (Kruskal-Wallis: p = 0.043), with *O. tauri* (median: 1.17, IQR=1.06-1.23) showing a trend for a moderately higher peak rate than *O. mediterraneus* (median: 0.94, IQR=0.83-0.98). Post-hoc Dunn tests did not identify significant pairwise differences after Holm correction (adj. p= 0.100), suggesting moderate but not strongly distinct group differences. Peak rate of the *O. mediterraneus* variant fell in an intermediate range (median: 1.06, IQR=1.02-1.09).

The thermal safety margin was similar across all species (Fig. 3e; Kruskal-Wallis: p=0.506; *O. mediterraneus* median: 4.01°C, IQR: 3.49-5.78°C; *O. mediterraneus* variant median: 3.60°C, IQR: 3.36-4.55°C; *O. tauri* median: 3.63°C, IQR: 2.91-4.35°C). Similarly, E_a_ showed no significant differences between species (ANOVA: F_(2, 22)_ = 0.667, p= 0.523). These two parameters, however, showed a high intraspecific variability (Fig 4a), with coefficients of variation (CVs) of species ranging from 28.4% to 45.9% for TSM and between 41.5% and 55.7% for E_a_. The most pronounced variability can be seen in *O. mediterraneus* (Fig. 4a). For T_opt_ interspecific variability was low, with CVs ranging from 5.1% to 6.5%. For peak rate, the *O. mediterraneus* variant markedly has the lowest within species-variation with a CV of 5.8%, showing a narrow range of peak rates (Fig. 4a), whereas variability was moderate within *O. tauri* (16.1%) and *O. mediterraneus* (19.1%).

**Figure 4.**
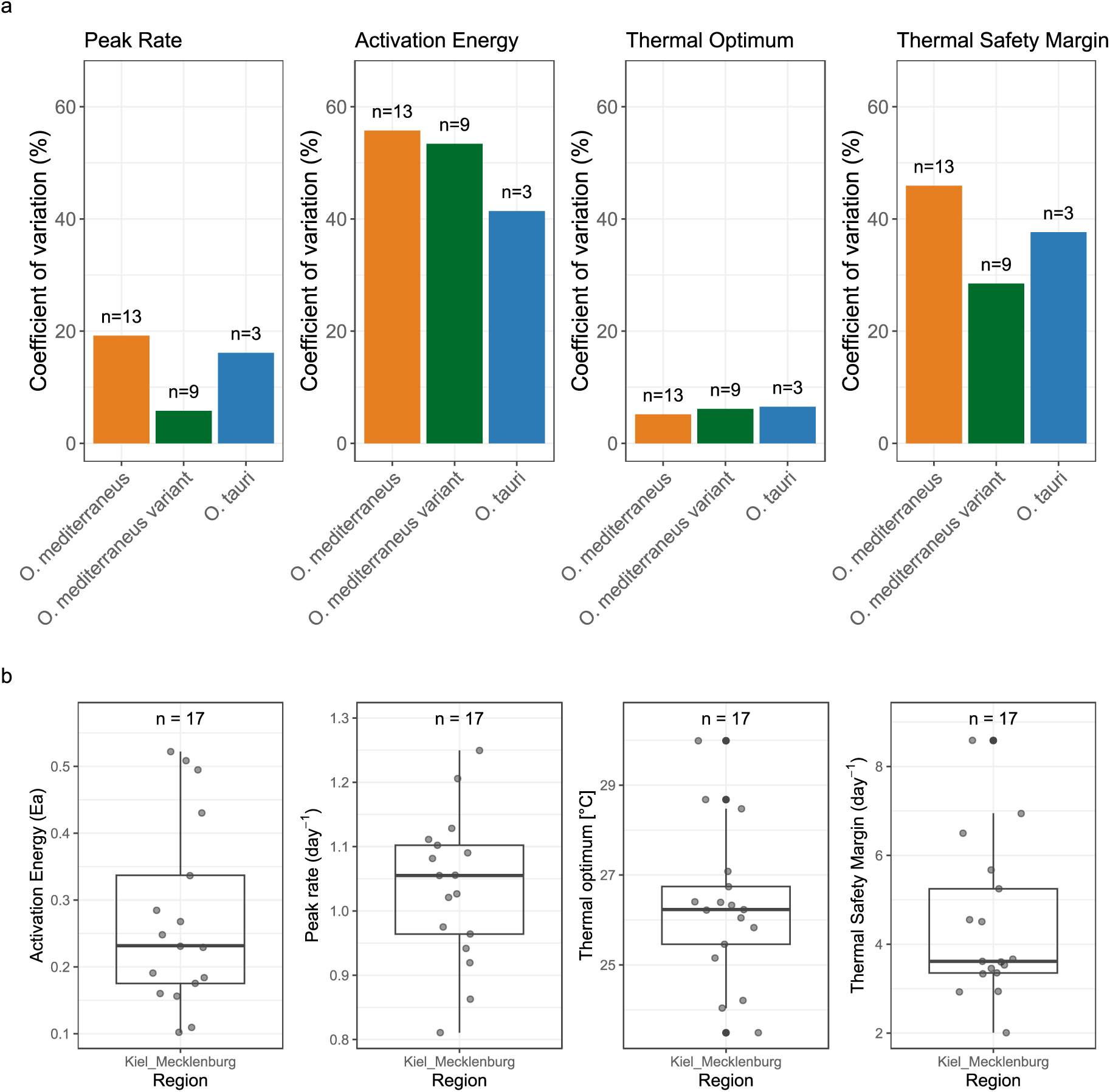
**a)** Intraspecific variability of thermal performance parameters peak rate, activation energy, thermal optimum and thermal safety margin displayed through the coefficient of variation for each species. The colors represent each species with orange for *O. mediterraneus*, green for the *O. mediterraneus* variant and blue for *O. tauri*. **b)** Variability of parameter estimates of strains originating from the Kiel region. The boxes of the boxplots represent the 25th to 75th percentile of the data with the line within the box representing the median (50th percentile), the whiskers include the smallest and largest values without the outliers (outside 1.5 × IQR; shown as points).

We also assessed intraregional variability within the thermally unpredictable (Santelia et al., 2026) Kiel area. For the 17 out of 27 strains that originated from this area, we examined the range and CVs of parameter values across strains. Parameter distributions spanned wide ranges across strains (Figure 4b). Similar to within species, T_opt_ and peak rate had lower intraregional variability (Supplemetary Figure S5) with T_opt_ ranging from 23.49°C to 29.99°C (median: 26.2°C, IQR: 25.5-26.7°C) and peak rate ranging from 0.81 d^−1^ to 1.24 d^−1^ (median: 1.06 d^−1^, IQR:0.96-1.10 d^−1^). E_a_ and TSM had higher intraregional variability (Supplementary Fig. S5) with E_a_ ranging from 0.10 to 0.52 (median:0.23, IQR: 0.18-0.34) and TSM ranging from 2.01°C to 8.56°C (median: 3.61°C, IQR:3.36-5.25°C).

### Species-specific temperature responses of size and chlorophyll a content across temperature

At baseline temperature (18°C, common garden treatment), size (ANOVA: F_(2, 90)_ = 7.260, p= 0.001) and chlorophyll *a* content proxy (Kruskal-Wallis: p= 0.001) differed significantly between species (Fig. 5a,b). Post-hoc tests revealed that for size and chlorophyll *a* content, *O. mediterraneus* significantly differed from both *O. tauri* (Size: Tukey’s HSD: p=0.012; Chl*a*: Dunn’s test: p=0.004) and the *O. mediterraneus* variant (Tukey’s HSD: p=0.006; Chl*a*: Dunn’s test: p=0.006).

**Figure 5.**
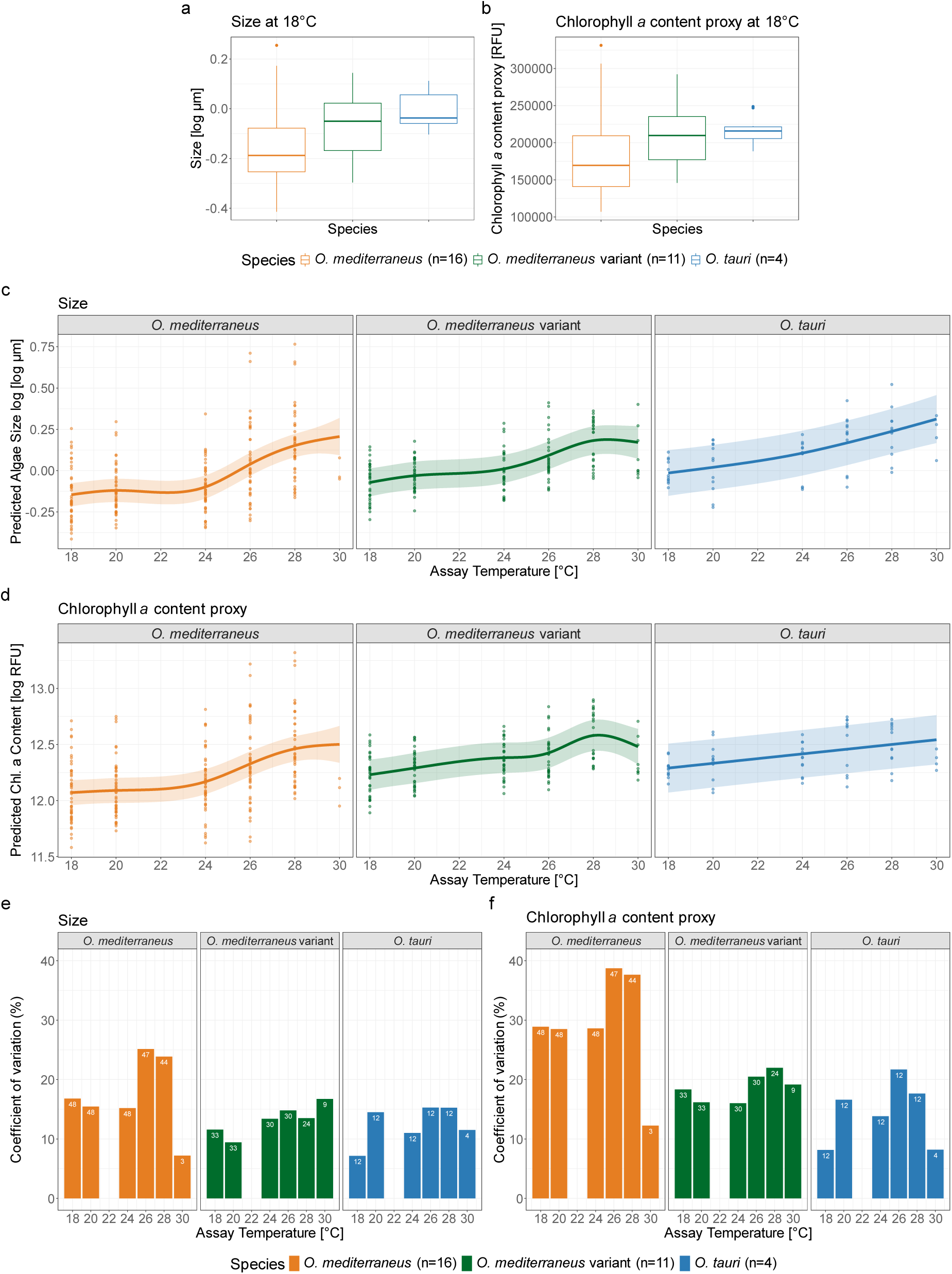
Species specific differences in **a)** size at 18°C at time of µmax (log size in µm) and **b)** proxy for chlorophyll a content (RFU) at 18°C at time of µmax. The boxes of the boxplots represent the 25th to 75th percentile of the data with the line within the box representing the median (50th percentile), the whiskers include the smallest and largest values without the outliers (outside 1.5 × IQR; shown as points). Temperature response of **c)** log size (log µm) and **d)** the proxy for chlorophyll *a* content, with GAMM fits between the three *Ostreococcus* species. Chlorophyll *a* proxy and size data are from the time point where µmax was reached. Colors indicate species, the shaded area indicates upper and lower confidence intervals (95%). The bar plots show intraspecific variability in **c)** size and **d)** chlorophyll *a* proxy across temperatures expressed through the coefficient of variation (%) at the respective temperature. The white numbers on the bars indicate the specific n for calculation, as replicates were included individually and varied across temperatures. The colors represent each species with orange for *O. mediterraneus*, green for the *O. mediterraneus* variant and blue for *O. tauri*.

Temperature had a significant effect on cell size in all three species (Fig. 5c; *O. mediterraneus*: edf = 4.21, F = 88.26, p < 0.001; *O. mediterraneus* variant: edf = 4.08, F = 36.80, p < 0.001; *O. tauri*: edf = 1.92, F = 45.08, p < 0.001). *O. tauri* displayed a more linear increase with warming, while O. *mediterraneus* and the *O. mediterraneus* variant had a lag at colder temperatures. *O. mediterraneus* showed a stronger slope of log size increase at warmer temperatures than the *O. mediterraneus* variant (Fig. 5a).

Temperature also had a significant effect on log chlorophyll *a* content across all species (Fig. 5d; *O. mediterraneus*: edf = 3.84, F = 70.65, p < 0.001; *O. mediterraneus* variant: edf = 4.51, F = 22.01, p < 0.001; *O. tauri*: edf = 1, F = 22.34, p < 0.001), with both *O. mediterraneus* and the *O. mediterraneus* variant showing non-linear responses, whereas *O. tauri* had a strong linear response. *O. mediterraneus* showed a pattern similar to that observed for cell size, with a lag at lower temperatures followed by an increase from 24 °C onwards. In contrast, the *O. mediterraneus* variant exhibited a more gradual increase up to 26 °C, followed by a steeper rise towards 28 °C and a decline at 30 °C. Interestingly, the temperature response of size and chlorophyll *a* of *O. tauri* was in line with the maximum growth rate, which increased linearly up to T_opt_, while for the other two species it was mostly decoupled from the maximum growth rate trend (Supplementary Fig. S6).

Intraspecific variability was moderate and tended to increase at warmer treatments (Fig. 5e,f) for both size and chlorophyll a content across all species. Notably, *O. mediterraneus* showed the highest variability within species, especially at 26°C and 28°C, both for log size and log chlorophyll *a* content with CVs reaching 25.1% for size and 38.7% for chlorophyl *a* content. Compared to *O. mediterraneus*, *O. tauri* and the O*. mediterraneus* variant only showed small changes in intraspecific variability between temperatures.

## Discussion

Assessing growth, metabolic and other trait variation within and across *Ostreococcus* species of the Baltic Sea improves our understanding of the evolutionary potential and ecological resilience of picophytoplankton under climate change. Our data revealed differing levels of variability, depending on the trait and species, suggesting contrasting degrees of physiological conservation, phenotypic plasticity, and adaptive potential among species.

The similarity in E_a_ and thermal safety margins among *Ostreococcus* species likely reflects conserved physiological constraints (Dell et al., 2011) combined with shared phylogeny (Barton & Yvon-Durocher, 2019), causing reduced interspecific variation. In addition, the similarity could also reflect adaptation to common environmental conditions, as Kontopoulos et al. (2020) found a systematic relationship between latitude and E_a_. In contrast to Barton and Yvon-Durocher (2019), who assessed one strain per species and reported high interspecific variability in E_a_, we observed substantial intraspecific variability, suggesting that species means derived from multiple strains may be more similar than estimates based on single strains.

In contrast, peak rate and T_opt_ showed significant interspecific differences but comparatively low intraspecific variability. These parameters are often shaped by ecological context, such as habitat temperature regimes or nutrient availability (Dell et al., 2011; Pang et al., 2024; M. K. Thomas et al., 2012) and thus might respond more quickly through both short-term acclimation and long-term selection in changing environments (Jin & Agustí, 2018; Listmann et al., 2016; Schaum et al., 2022). Schaum et al. (2022) demonstrated that especially in fluctuating environments both T_opt_ and peak rate have the potential to evolve rapidly. The strong environmental variability of the Baltic Sea, driven by its unique topography, anthropogenic influences, increasing heatwave events, and pronounced daily summer temperature fluctuations (Frölicher et al., 2018; Lindenthal et al., 2024; Paalme et al., 2020; Pinto et al., 2024), may therefore promote divergence in T_opt_ and peak rate between species, while remaining more conserved within species.

Interspecific variability in T_opt_ and peak rate may allow for niche complementarity and partitioning, where different species perform optimally at different temperatures, reducing direct competition and enhancing overall community stability and productivity across thermal gradients (Godoy et al., 2020; Litchman et al., 2012; Replansky & Bell, 2009; M. K. Thomas et al., 2012; Vallina et al., 2017). This functional redundancy and diversity align with the “insurance hypothesis” (McNaughton, 1977; Yachi & Loreau, 1999), which argues that biodiversity increases resilience against environmental change. Interspecific differences in T_opt_ may also influence seasonal succession, where species with lower T_opt_ may dominate in spring and fall, while warm-adapted species could succeed in summer months. Santelia et al. (2022) found seasonality to be the main driver of differences in the T_opt_ of gross photosynthesis for Baltic Sea phytoplankton communities and, Alegria Zufia et al. (2021) reported distinct seasonal and thermal ranges for different picophytoplankton groups.

The evolution of specific thermal traits relies on within species selection either when new mutations occur or via sorting of genotypes (Costas et al., 2014; Listmann et al., 2016). High intraspecific variability in thermal responses, as observed for E_a_, TSM, size and chlorophyll *a* content in all of our study species, may be an important trait variation for selection to act on. It can facilitate buffering of populations against environmental fluctuations, particularly in the face of ongoing climate change (Kremp et al., 2012). Intraspecific variability provides a form of diversified bet-hedging, whereby different phenotypes within a population are optimized for slightly different thermal conditions, increasing the likelihood that some will succeed as conditions change (Cohen, 1966; Kling et al., 2023; Philippi & Seger, 1989; Seger & Brockmann, 1987). Intraspecific trait variability has not only been shown between strains from different isolation times and locations (Krinos et al., 2025), but also among coexisting strains (Ajani et al., 2021). Sjöqvist and Kremp (2016) demonstrated that intraspecific diversity can be beneficial under salinity stress, indicating a stabilizing impact of increased diversity. The high intraregional variability of strains within the Kiel region further aligns with findings by Bishop et al. (2022), who found high thermal trait variation for Southern Ocean diatoms. Furthermore, Santelia et al. (2026) showed that that the Kiel area is characterized by a high thermal variability and that this lead to a higher degree of phenotypic plasticity for phytoplankton from that area. These findings support the idea that intraspecific diversity is a key ecological strategy in variable environmental conditions.

While all species exhibit similar curve width and thermal safety margins across the tested temperatures, they may achieve this through different evolutionary strategies. Especially *O. mediterraneus* exhibited consistently high intraspecific variability in thermal traits and curves, accompanied by high variability and the most prominent changes in the temperature response of chlorophyll *a* and size. In addition, *O. mediterraneus* showed decoupled thermal responses of size, chlorophyll *a* content, and growth rate, with increases in size and chlorophyll *a* despite declining growth, possibly reflecting increased energetic allocation to maintenance and repair under thermal stress (Leles & Levine, 2023). Together, these patterns suggest a high degree of phenotypic plasticity in *O. mediterraneus*, enabling flexible physiological responses to changing thermal conditions.

Contrary to *O. mediterraneus*, *O. tauri* exhibited consistent linear increases of size, chlorophyll *a* content and growth rate with warming, which suggests a coordinated physiological adjustment with little trade-offs across temperature, possibly highlighting a broader thermal tolerance and lower phenotypic plasticity. Consistent with this interpretation, *O. tauri* strains exhibiting some of the lowest intraspecific variability in size and chlorophyll *a* content and the lowest range of T_opt_ and E_a_ values among the three species. Although this reduced variability could partially result from the smaller sample size for *O. tauri*, the analyzed strains originated from geographically diverse sampling stations. Generally, evidence suggests *O. tauri* has a broader global distribution than *O. mediterraneus* (Demir-Hilton et al., 2011) and is present in more locations within the Baltic Sea (Hu et al., 2016). Therefore, it might have evolved a robust generalist strategy which often is characterized by broad but flat thermal performance curves, which may reduce sensitivity to environmental variation (Kaspari et al., 2016).

Similar diversification of thermal strategies as seen between *O. mediterraneus* and *O. tauri* has been observed in diatoms, where species adapt either through shifts in thermal optima or through a switch from specialist to generalist, expanding their maximum critical thermal limit (Jin & Agustí, 2018). This shows that different strategies can be successful. Variation in phenotypic plasticity could further support *Ostreococcus* populations persistence and adaptation (Walter et al., 2023), especially under directional selection pressures such as climate impacts.

## Conclusion

Overall, our findings highlight the potentially adaptive and stabilizing role of inter- and especially intraspecific variation of traits in response to temperature. In the Baltic Sea *Ostreococcus* species system, we see a population level diversification of bet-hedging with high phenotypic plasticity and high intraspecific variability combined with more generalist phenotypes with lower phenotypic plasticity and intraspecific variability. Together, these strategies can enhance community-level stability, ensuring that *Ostreococcus* contributes to consistent primary production and carbon cycling despite both short-term variability and longer-term shifts in temperature regimes. In addition, the variability within the Kiel area further suggests that intraregional variability could be an ecological strategy for persistence in variable and changing environments. Future studies aiming to model microbial responses to climate change or construct predictive thermal niche models should account for not only species-level averages but also the variance within species and regions (Bennett et al., 2019), as this variability can fundamentally alter predictions of resilience, range shifts, and ecosystem function (Bishop et al., 2022; Valladares et al., 2014; Wolf et al., 2018).

## Author contributions

ES and LL conceived the study. CP and MV designed and conducted the experiments, analyzed the data, produced the figures and wrote the first draft of the manuscript. CP wrote the final draft of the manuscript. ES contributed to coding and code troubleshooting. ES and LL contributed with analysis input and revisions of the manuscript. All authors gave final approval for publication. Funding was acquired by ES and LL.

## Acknowledgements

We want to thank our lab technician Stefanie Schnell and the student Janice da Mata for support during the experiment. This research was supported through funding of the Deutsche Forschungsgemeinschaft DFG grant 471040156. ChatGPT (OpenAI, GPT-5.1) was used for language and grammar improvement, it was operated in a mode where user inputs were not used for model training and all outputs were reviewed and validated by the authors.

## Data Availability

The data supporting the findings of this study are available in the Supporting Information. R codes and data sets used for analysis are available on Zenodo (10.5281/zenodo.20430934).

## Abbreviations

Chl*a*: Chlorophyll *a* content proxy
T_opt_: Thermal optimum
TSM: Thermal safety margin
E_a_: Activation Energy
µmax: Maximum growth rate
CV: coefficient of variation
*O. tauri*: *Ostreococcus tauri*
*O. mediterraneus*: *Ostreococcus mediterraneus*
*O. mediterraneus* variant: *Ostreococcus mediterraneus* variant

