## SupplementaryMaterial for "Intraspecific variability exceeds interspecific variability in thermal responses of Baltic Sea *Ostreococcus* (Mamiellophyceae) traits"

**For**

**'Shared last author**

[1]Institute of Marine Ecosystem and Fisheries Science, University of Hamburg, Hamburg, Germany

### Supplementary Figures

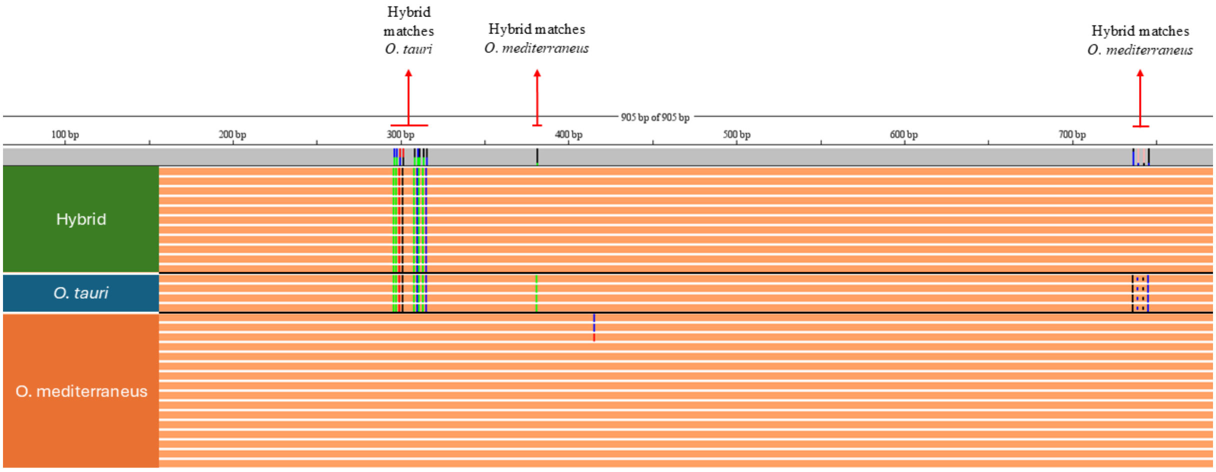


**Supplementary Fig. S1** Annotated screenshot of the 18S alignment in Codon Code Aligner, outlining the difference in 18S sequences of the hybrid strains in relation to *Ostreococcus tauri* and *Ostreococcus mediterraneus* strains. Red arrows point to the relevant sequence parts.

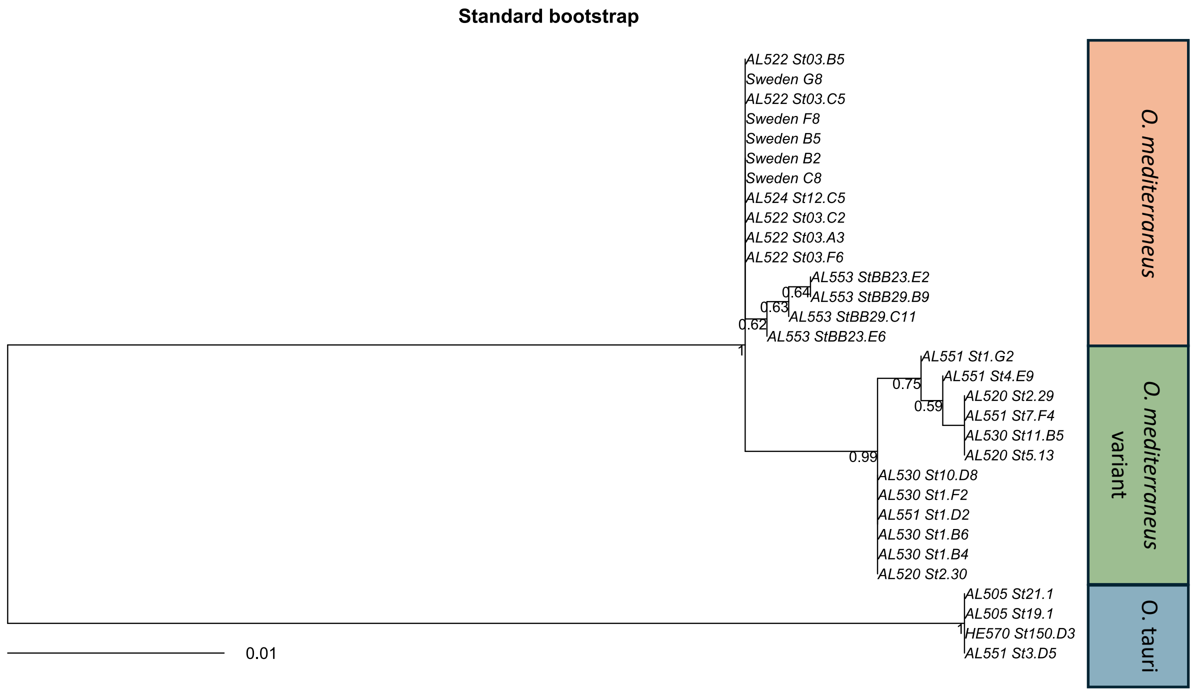


**Supplementary Fig. S2** Maximum likelihood phylogenetic tree based on ITS-region sequences (1018 bp) of the 31 *Ostreococcus* strains used in this study. The best-fitting nucleotide substitution model (JC) was selected using model testing (BIC). Numbers at the nodes indicate standard bootstrap support values (1000 replicates). Boxes on the right-hand side denote species-level assignments of the strains.

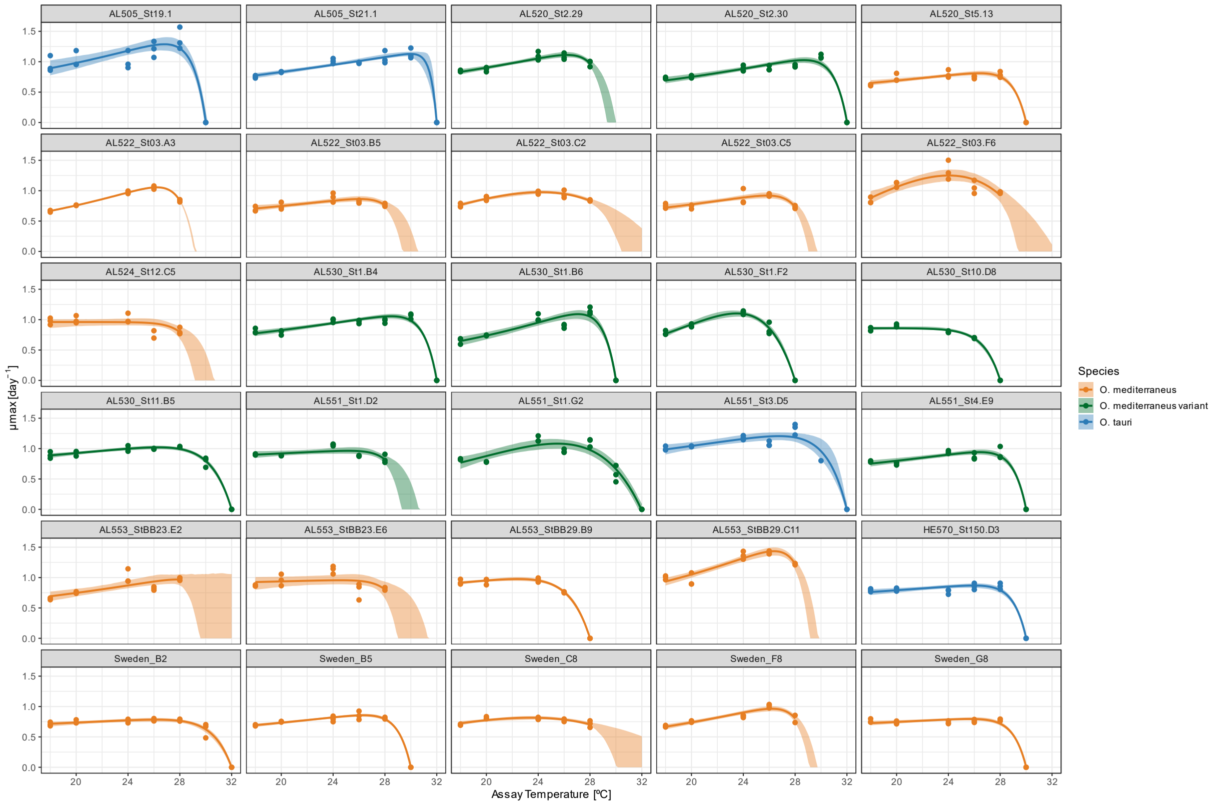


**Supplementary Fig. S3 Fitted thermal performance curves per strain.** Predicted model fits (lines) and extracted μmax data points (points) are shown across assay temperatures. Shaded areas show 95% confidence intervals from bootstrapping (1000 iterations). Colors represent the different species, with orange showing *O. mediterraneus*, green showing the *O. mediterraneus* variant and blue showing *O. tauri*.

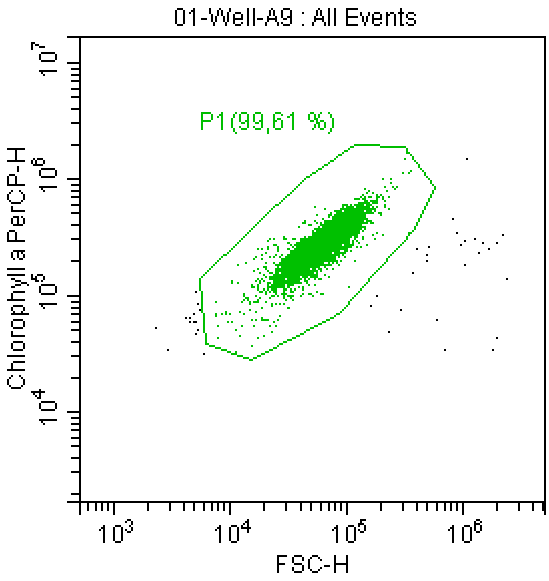


**Supplementary Fig. S4:** Example of flow cytometry gating used to define phytoplankton populations for cell counts. Fluorescence and forward scatter parameters were used to distinguish phytoplankton cells from background noise and debris. The gated region (outlined in green represents the population included in downstream cell count analyses.

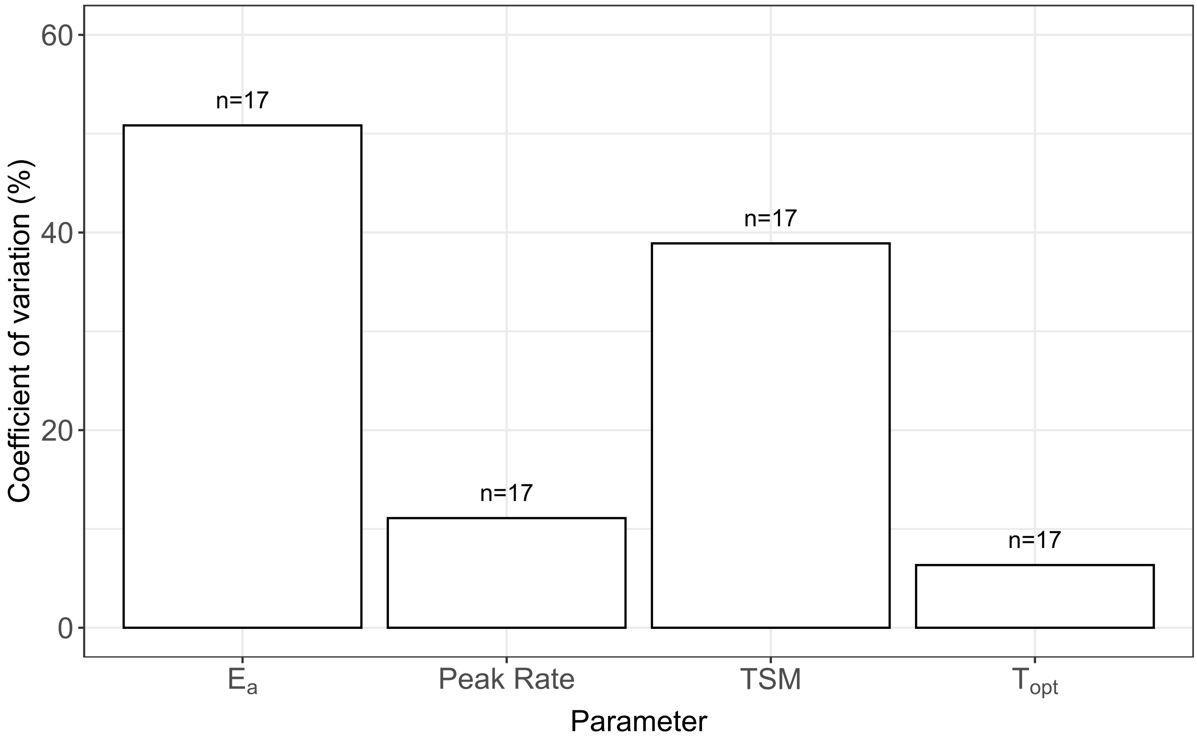


**Supplementary Fig. S5** Coefficients of variation (%) for the thermal performance parameters activation energy (Ea), peak rate, thermal safety margin (TSM) and thermal optimum (Topt) for the 17 strains from the Kiel-Mecklenburg area. Coefficients of variation were calculated as the standard deviation divided by the mean for each parameter across strains.

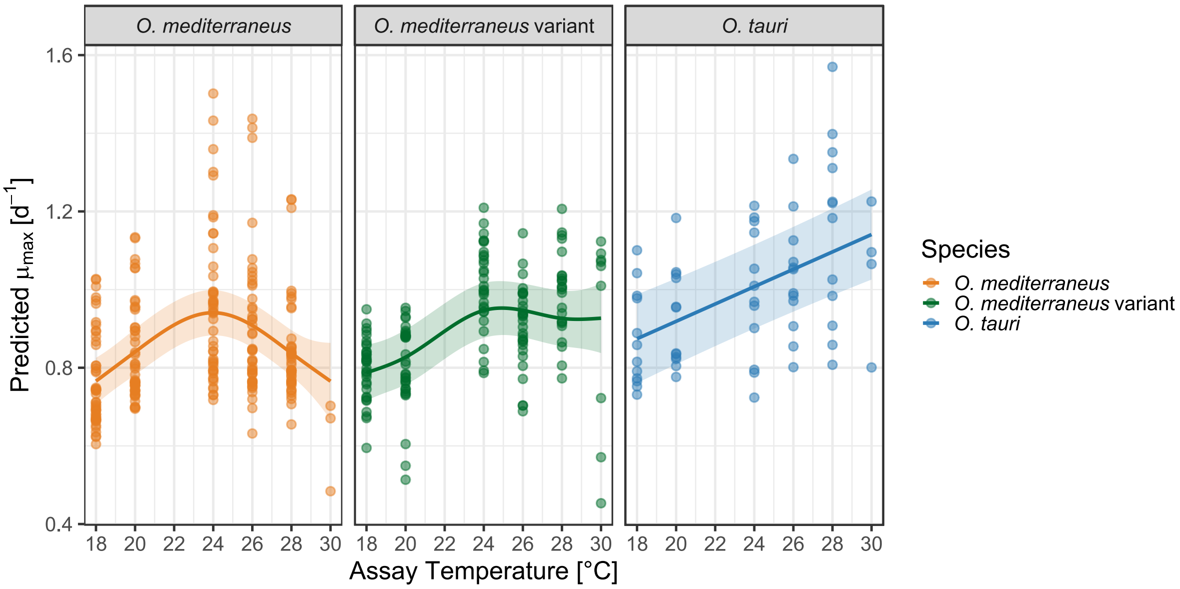


**Supplementary Fig. S6** Species specific differences in mumax $d^{-1}$ (points) across temperature with GAMM fits (lines) between the three *Ostreococcus* species. Colors indicate species, the shaded area indicates upper and lower confidence intervals (95%) of GAMM fits.

### Supplementary Tables

**Supplementary Table 1** Overview of which strain was grown and measured at what temperature during which Batch. Temperatures written in red were excluded due to light issues within the incubators. Between batch 1 and 2 the 22°C degree samples were used as a repeating control between batches. This is still valid as temperature was constant at the 22°C treatment but much lower than the other temperatures (and therefore excluded). As a control between Batch 2 and 3 the 18°C treatment was used as a repeating control. White boxes indicate it was not tested at that temperature treatment.

| | Batch 1 | | | | Batch 2 | | | | | Batch 3 | | | | | |
|---|---|---|---|---|---|---|---|---|---|---|---|---|---|---|---|
| **Strain** | 22°C | 24°C | 26°C | 28°C | 15°C | 18°C | 20°C | 22°C | 26°C | 15°C | 18°C | 22°C | 28°C | 30°C | 32°C |
| AL505_St19.1 | | | | | | | | | | | | | | | |
| AL505_St21.1 | | | | | | | | | | | | | | | |
| AL520_St5.13 | | | | | | | | | | | | | | | |
| AL520_St2.29 | | | | | | | | | | | | | | | |
| AL520_St2.30 | | | | | | | | | | | | | | | |
| AL522_St03.A3 | | | | | | | | | | | | | | | |
| AL522_St03.B5 | | | | | | | | | | | | | | | |
| AL522_St03.C2 | | | | | | | | | | | | | | | |
| AL522_St03.C5 | | | | | | | | | | | | | | | |
| AL522_St03.F6 | | | | | | | | | | | | | | | |
| AL524_St12.C5 | | | | | | | | | | | | | | | |
| AL530_St1.B4 | | | | | | | | | | | | | | | |
| AL530_St1.B6 | | | | | | | | | | | | | | | |
| AL530_St1.F2 | | | | | | | | | | | | | | | |
| AL530_St10.D8 | | | | | | | | | | | | | | | |
| AL530_St11.B5 | | | | | | | | | | | | | | | |
| Sweden_B2 | | | | | | | | | | | | | | | |
| Sweden_B5 | | | | | | | | | | | | | | | |
| Sweden_C8 | | | | | | | | | | | | | | | |
| Sweden_F8 | | | | | | | | | | | | | | | |
| Sweden_G8 | | | | | | | | | | | | | | | |
| HE570_St150.D3 | | | | | | | | | | | | | | | |
| AL551_St1.D2 | | | | | | | | | | | | | | | |
| AL551_St1.G2 | | | | | | | | | | | | | | | |
| AL551_St3.D5 | | | | | | | | | | | | | | | |
| AL551_St4.E9 | | | | | | | | | | | | | | | |
| AL551_St7.F4 | | | | | | | | | | | | | | | |
| AL553_StBB23.E2 | | | | | | | | | | | | | | | |
| AL553_StBB23.E6 | | | | | | | | | | | | | | | |
| AL553_StBB29.B9 | | | | | | | | | | | | | | | |
| AL553_StBB29.C11 | | | | | | | | | | | | | | | |

**Supplementary Table 2** Initial model parameters for nlsLoop modified gompeortz model growth curve fitting. LOG10N0 = starting cells as log10, LOG10MAX= carrying capacity as log10, mumax= µmax, lag = lag phase length in days, model was tried 1000 times.

| Parameter | Starting value lower | Starting value upper |
|---|---|---|
| LOG10N0 | 2 | 6 |
| LOG10Nmax | 4 | 10 |
| mumax | 0.1 | 2.5 |
| lag | 0.001 | 12 |

**Supplementary Table 3** Model parameter estimates and 95% confidence intervals (in brackets) from bootstrapping (1000 iterations) per strain. * indicates strains where all parameters were excluded from further analysis ** indicates strains where only $E_a$ was excluded from further analysis.

| Strain | Model | Peak rate | $T_{opt}$ [°C] | $E_a$ | TSM [°C] |
|---|---|---|---|---|---|
| AL505_St19.1 | tomlinson | 1.29 (1.18-1.41) | 26.81 (26.34-27.06) | 0.1805 (0.1884-0.4750) | 3.21 (2.83-3.68) |
| AL505_St21.1 | lactin2 | 1.13 (1.09-1.18) | 29.99 (29.03-31.29) | 0.2474 (0.2020-0.3091) | 2.01 (0.71-2.97) |
| AL520_St2.29 | tomlinson | 1.11 (1.07-1.16) | 26.23 (26.06-26.48) | 0.2684 (0.2120-0.3446) | 3.36 (3.11-3.54) |
| AL520_St2.30 | tomlinson | 1.03 (0.98-1.08) | 28.68 (28.46-28.89) | 0.1915 (0.2257-0.3631) | 3.33 (3.11-3.54) |
| AL520_St5.13 | tomlinson | 0.81 (0.78-0.85) | 26.39 (25.92-26.63) | 0.1601 (0.1323-0.2805) | 3.61 (3.25-3.96) |
| AL522_St03.A3 | tomlinson | 1.06 (1.04-1.07) | 26.22 (26.2-26.2) | 0.4309 (0.4186-0.4695) | 2.94 (2.83-2.97) |
| AL522_St03.B5 | tomlinson | 0.86 (0.83-0.91) | 26.05 (25.64-26.63) | 0.1839 (0.1109-0.2794) | 3.66 (3.25-4.1) |
| AL522_St03.C2 | lactin2 | 0.98 (0.95-1) | 24.21 (23.94-25.35) | 0.2847 (0.2167-0.3526) | 8.58 (4.94-8.06) |
| AL522_St03.C5 | tomlinson | 0.92 (0.88-0.97) | 25.83 (25.49-26.2) | 0.2316 (0.1519-0.3436) | 3.46 (3.11-3.82) |
| AL522_St03.F6 | lactin2 | 1.25 (1.16-1.34) | 24.04 (23.66-25.49) | 0.5093 (0.2438-0.5949) | 6.95 (3.82-7.78) |
| AL524_St12.C5* | tomlinson | 0.96 | 18.00 | NA | 11.76 |
| AL530_St1.B4 | tomlinson | 1.06 (1.02-1.1) | 28.47 (28.18-28.61) | 0.1752 (0.1821-0.2914) | 3.53 (3.25-3.68) |
| AL530_St1.B6 | tomlinson | 1.09 (1.02-1.15) | 27.08 (26.77-27.33) | 0.3369 (0.3432-0.5758) | 2.93 (2.69-3.11) |
| AL530_St1.F2 | lactin2 | 1.10 (1.07-1.15) | 23.49 (23.09-23.8) | 0.5220 (0.4298-0.6219) | 4.51 (4.1-4.81) |
| AL530_St10.D8* | lactin2 | 0.86 | 18.00 | NA | 10.00 |
| AL530_St11.B5 | lactin2 | 1.02 (0.99-1.05) | 26.33 (25.78-26.77) | 0.1098 (0.0840-0.1853) | 5.67 (5.09-6.22) |
| AL551_St1.D2 | tomlinson | 0.96 (0.92-1.02) | 25.15 (18-25.92 | 0.2295 (0.0080-0.1759) | 4.55 (3.68 -11.88) |

| | | | | | |
|---|---|---|---|---|---|
| AL551_St1.G2 | lactin2 | 1.08 (1.01-1.16) | 25.46 (24.51-26.2) | 0.4955 (0.2011-0.5413) | 6.50 (5.66-7.49) |
| AL551_St3.D5 | lactin2 | 1.21 (1.15-1.28) | 26.74 (25.49-28.47) | 0.1014 (0.0872-0.2947) | 5.25 (3.39-6.36) |
| AL551_St4.E9 | tomlinson | 0.94 (0.9-0.99) | 26.40 (26.06-26.63) | 0.1563 (0.1419-0.2911) | 3.60 (3.25-3.96) |
| AL553_StBB23.E2* | tomlinson | 0.97 | 27.76 | 0.2649 | 3.39 |
| AL553_StBB23.E6 | tomlinson | 0.96 (0.88-1.06) | 24.53 (18-26.2) | 0.3216 (0.0038-0.2345) | 5.18 (3.39-13.01) |
| AL553_StBB29.B9 | lactin2 | 0.98 (0.99-1.05) | 22.41 (22.1-22.81) | 0.0379 (0.1598-0.2972) | 5.59 (5.09-5.94) |
| AL553_StBB29.C11 | tomlinson | 1.43 (1.37-1.49) | 26.37 (26.2-26.63) | 0.3975 (0.3187-0.4911) | 3.02 (2.83-3.11) |
| HE570_St150.D3** | tomlinson | 0.87 (0.83-0.91) | 25.96 (25.35-26.34) | -0.0408 | 4.05 (3.54-4.67) |
| Sweden_B2 | lactin2 | 0.78 (0.75-0.81) | 26.16 (25.21-26.91) | 0.0749 (0.0301-0.1611) | 5.84 (5.09-6.79) |
| Sweden_B5 | tomlinson | 0.86 (0.83-0.88) | 26.35 (26.2-26.48) | 0.1776 (0.1596-0.2439) | 3.65 (3.39-3.82) |
| Sweden_C8 | lactin2 | 0.82 (0.79-0.84) | 23.42 (22.81-25.21) | 0.5451 (0.0717-0.2275) | 10.67 (4.95-9.19) |
| Sweden_F8 | tomlinson | 0.96 (0.92-1) | 26.22 (25.92-26.48) | 0.3526 (0.2708-0.4298) | 3.14 (2.83-3.25) |
| Sweden_G8** | tomlinson | 0.79 (0.77-0.82) | 25.65 (24.65-26.06) | -0.0343 | 4.36 (3.82-5.23) |

**Supplementary Table 4** Boundaries and settings for calculating the parameters from the modelfits (function: "calc_params()") within the rTPC package (Daniel Padfield et al. 2026). Bootstrap values were calculated following the same methods.

| **Parameter** | **Boundaries/Settings** |
|---|---|
| **peak rate ("rmax" in rTPC package)** | Picks the maximum rate value of the predictions within the temperature range (default of rTPC package) |
| **$T_{opt}$ ("topt" in rTPC package) in °C** | Picks the temperature with the largest rate value occurring, accurate up to 0.001°C (default of rTPC package) |
| **$E_a$ ("e" in rTPC package)** | Fits a modified-Boltzmann equation to all raw data below Topt (default of rTPC package) |
| **thermal safety margin ("thermal_safety_margin" in rTPC package)** | Thermal safety margin is calculated as: CTmax – Topt (default of rTPC package) |
